# The relationship between body size and skull and dental characters in cricetid rodents from Nebraska

**DOI:** 10.64898/2025.12.12.693729

**Authors:** Opal K. A. L. Wagner, S. Kathleen Lyons

## Abstract

Body size in mammals spans 20 orders of magnitude and is strongly correlated with ecology, making it one of the most important characteristics for a mammal. Body mass must be estimated for fossil mammals because the whole organism is rarely fully preserved. Although the relationships between anatomical characters and body mass have been calculated for many groups of mammals, rodents are less well studied. Previous work had limited sampling of species or used a species average for body size and may not be accurate. Because rodents are the most species rich order of mammals having accurate relationships for the different families of rodents is crucial for understanding the evolution of body size in rodents. I analysed the relationship between body size and three skull characters (skull, molar (M1), and toothrow length) in 11 species of cricetid rodents from Nebraska using reduced major axis regressions. I found a significant positive relationship between all characters and body size. Skull and toothrow length are the best predictors of body size. M1 length is not as good of a predictor because one genus (*Microtus*) has an unusually large M1 length for its body size. Separate regression equations for the Arvicolines and the Sigmodontids provide better predictors for body size using lower M1 length. This work will allow for more accurate predictions of body mass in extinct cricetids.

## Introduction

Body size is one of the most important characteristics of a mammal. Body size in mammals spans 20 orders of magnitude, from the size of a shrew to the size of a blue whale (Smith and Lyons 2013). The size of a mammal species is strongly correlated with its ecology (Brown 1995, Brown, et al. 2004, Brown and Sibly 2006, Calder 2000, 1983, Damuth 1981, 1981, Ernest, et al. 2003, Kelt and Van Vuren 1999, Peters 1983, Smith and Lyons 2013, 2011, Smith, et al. 2008). Mammals of different sizes have different life spans, diets, and reproduction. For example, larger mammals live longer than smaller mammals, and are more likely to be herbivores (Brown and Sibly 2006, Ernest 2003, Peters 1983, Sibly and Brown 2007). In addition, small mammals have faster reproduction because they have larger litter sizes, and shorter intervals between reproduction. Therefore, being able to estimate body size in a mammal species tells you a lot about its lifestyle.

Body mass estimation in fossils is difficult due to the fact that the whole organism is rarely fully preserved. In particular, mammal fossils consist mostly of isolated teeth, jaws, and skulls. The rest of the skeletal structure is rarely preserved (Behrensmeyer 1984). Fortunately, the relationships between body size and anatomical characters for modern organisms can be used to estimate body size in extinct animals. For example, Van Valkenburgh (1990) found that molar length in modern carnivores is strongly related to body mass. Van Valkenburgh’s (1990) regression equations are commonly used to estimate the mass of extinct carnivores e.g, (Smith, et al. 2022).

Although the relationships between anatomical characters and body mass have been calculated for many groups of mammals (e.g., Damuth and MacFadden 1990), rodents are less well studied. Studies that have been done have had limited sampling of species and did not include Cricetidae (Bertrand, et al. 2015), or used a species average for body size instead of measurements from individual specimens (Freudenthal and Martín-Suárez 2013), and thus may not be accurate. Because rodents are the most species rich order of mammals and make up a majority of the mammal fossil record in North America (Alroy 1998), having accurate relationships for the different families of rodents is crucial for understanding the evolution of body size in rodents.

The Cricetidae is a diverse family within the order Rodentia that has over 800 species world-wide, 16 of which occur in Nebraska. Two subfamilies, the Arvicolinae and Sigmodontinae, occur throughout North America in almost all terrestrial habitats including the arctic tundra, forests and grasslands. Cricetids fill many different niches; some are terrestrial, some are arboreal and at least one genus, *Ondatra*, the muskrat, is semi-aquatic. Their diets range from herbivorous to insectivorous, with a few species found in Nebraska being carnivorous (e.g., *Onychomys*, the grasshopper mouse). The species found in Nebraska encompass the full range of body sizes found in this family from the tiny genus (∼10 g), *Reithrodontomys* to the much larger genus (∼1 kg), *Ondatra* (Wilson and Reeder 2005). Thus, this is an ideal family to explore the relationship between skull and dental characters and body size.

Here, I quantify the relationship between body size and skull length, lower tooth row length, and lower first molar length in cricetid rodents from Nebraska. My goal was to determine the best relationship for estimating body size in extinct Cricetidae. Thus, I asked 1: what is the relationship between body size and skull length, toothrow length, and lower first molar length in cricetids? 2: Which skull or dental character has the strongest relationship with body size and thus is the best for estimating body size in fossil cricetids? 3: Is the relationship the same for the subfamilies Arvicolinae and Sigmodontinae?

## Methods and Materials

### Data

The specimens used in this study are housed in the University of Nebraska State Museum (UNSMZ). I examined cranial and dental characters from 220 specimens across 11 species of Cricetidae that occur in Nebraska. Specimens with associated field weights were preferentially sampled. Thus, all specimens measured have a unique associated body mass. I sampled 10 males and 10 females from 10 of the 11 species. One species, *Neotoma cinerea*, only had 9 males with associated field weights. For that species, I measured 9 males and 11 females. For all specimens, I recorded museum number, taxonomic identity, sex, weight (g), skull length (mm), lower right first molar length (mm), and lower right toothrow length (mm). These skull and dental characters were chosen because they are commonly measured in studies that are trying to predict body mass using cranial and dental characters (Bertrand, et al. 2015, Freudenthal and Martín-Suárez 2013, Gingerich and Smith 2025, Hopkins 2018, Hopkins 2008, Martin 1993, Martin, et al. 2009, Millien and Bovy 2010) making this study directly comparable. These data are recorded in Table S1 in the Supplementary Materials.

### Analyses

First, I calculated the geometric mean of the body mass of males and females separately for each species. I compared the body mass of males and females for each species using t-tests to determine if any of the species were sexually dimorphic. Next, I calculated the geometric mean and standard deviation for all measured skull and dental characters to determine if the variation within each species was similar. Means and standard deviations for all measured skull and dental characters were calculated for males and females pooled because no species showed significant differences in body mass between males and females. Third, I used reduced major axis (RMA) regressions to determine the relationship between skull and dental characters and body mass for all 11 species as RMA regressions account for measurement error in both the independent and dependent variables. In this study the measurements of both body size and the skull and dental characters have errors associated with them. Finally, I used RMA regressions to analyze the relationship between skull length, lower right M1 length, lower right toothrow length and body size separately for the Arvicolinae and the Sigmodontinae. Body mass was logged in all analyses.

## Results

None of the species included in this study showed significant sexual dimorphism (Table 1). Therefore, data for males and females were pooled for all subsequent analyses. The standard deviations of each species body mass were similar suggesting that uncertainty in the body mass data is similar across all species (Table 2). Therefore, analysing the relationship between body size and skull and dental characters using reduced major axis regression on the raw data across species is reasonable.

**Table 1.** T-test results for sexual dimorphism in cricetid rodents. None of the included species show significant differences in body mass between males and females.

| Species | t Statistic | p-value | Geometric Mean Female Mass (g) | Geometric Mean Male Mass (g) | Standard Deviation Female Mass | Standard Deviation Male Mass |
| --- | --- | --- | --- | --- | --- | --- |
| *Microtus ochrogaster* | -1.08 | 0.30 | 43.77 | 37.93 | 0.84 | 0.96 |
| *Microtus pennsylvanicus* | 0.86 | 0.40 | 36.17 | 39.77 | 0.92 | 1.03 |
| *Ondatra zibethicus* | 0.24 | 0.81 | 825.49 | 850.57 | 0.96 | 0.88 |
| *Neotoma cinerea* | 0.30 | 0.77 | 192.28 | 199.74 | 0.94 | 0.89 |
| *Neotoma floridana* | 1.43 | 0.17 | 268.92 | 317.09 | 1.09 | 0.87 |
| *Onychomys leucogaster* | -0.21 | 0.83 | 31.41 | 30.67 | 0.87 | 1.13 |
| *Peromyscus leucopus* | -0.10 | 0.92 | 22.18 | 21.92 | 0.89 | 1.01 |
| *Peromyscus maniculatus* | 0.45 | 0.66 | 21.08 | 21.91 | 1.03 | 1.16 |
| *Reithrodontomys megalotis* | -1.02 | 0.32 | 10.67 | 9.39 | 0.89 | 0.94 |
| *Reithrodontomys montanus* | -1.75 | 0.10 | 10.56 | 9.11 | 1.05 | 1.13 |
| *Sigmodon hispidus* | -1.05 | 0.31 | 98.81 | 81.69 | 0.74 | 0.77 |

**Table 2.** Mean and standard deviation of all skull and dental characters for each species. Mean values of skull and dental characters differ among species, but the standard deviations are similar suggesting that variation within species is smaller than between species.

| Species | Geometric Mean Skull Length (mm) | Standard Deviation Skull Length (mm) | Geometric Mean Lower Right Toothrow Length (mm) | Standard Deviation Lower Right Toothrow Length (mm) | Geometric Mean Lower Right M1 Length (mm) | Standard Deviation Lower Right M1 Length (mm) |
| --- | --- | --- | --- | --- | --- | --- |
| *Microtus ochrogaster* | 26.85 | 1.07 | 6.23 | 1.08 | 3.23 | 1.08 |
| *Microtus pennsylvanicus* | 26.79 | 1.06 | 6.46 | 1.07 | 3.33 | 1.09 |
| *Ondatra zibethicus* | 60.02 | 1.05 | 14.28 | 1.05 | 7.48 | 1.06 |
| *Neotoma cinerea* | 44.08 | 1.08 | 8.78 | 1.04 | 3.46 | 1.08 |
| *Neotoma floridana* | 47.28 | 1.06 | 8.77 | 1.03 | 3.51 | 1.06 |
| *Onychomys leucogaster* | 25.99 | 1.05 | 4.23 | 1.07 | 1.86 | 1.08 |
| *Peromyscus leucopus* | 24.42 | 1.06 | 3.93 | 1.05 | 1.60 | 1.06 |
| *Peromyscus maniculatus* | 23.42 | 1.04 | 3.68 | 1.08 | 1.54 | 1.08 |
| *Reithrodontomys megalotis* | 18.73 | 1.06 | 3.07 | 1.03 | 1.39 | 1.09 |
| *Reithrodontomys montanus* | 18.31 | 1.04 | 2.30 | 1.04 | 1.38 | 1.05 |
| *Sigmodon hispidus* | 30.74 | 1.08 | 6.23 | 1.05 | 2.33 | 1.08 |

Skull length, lower right toothrow length, and lower right first molar length all had a significant relationship with body size (Table 3, Fig. 1). However, the predictive power of molar length is much less than either sull length or toothrow length when analysing the family as a whole. Inspection of the graphs (Fig. 1 C) suggests that the species of the genus Microtus have a different relationship between M1 length and body size. In particular they have large M1 for their body size in comparison with other members of the family. The relationships between lower right M1 length and body size and lower right toothrow length improved when the subfamilies were analysed separately (Table 4, Fig. 2). While the slope of the relationship between body size and lower right M1 was similar for Arvicolinae and Sigmodontinae, the intercepts were offset. For the relationship between body size and lower right toothrow length, the subfamilies had both different slopes and intercepts. Nonetheless, the relationship between body size and skull length was not appreciably different for the subfamilies than for the family as a whole (Tables 3, 4, Figs. 1, 2).

**Table 3.** Reduced major axis regressions for the relationship between skull and dental characters and body mass in cricetid rodents. Skull length (mm), toothrow length (mm), and lower first molar length are all significantly related to body mass in cricetid rodents. The *R*^2^ for skull length is the highest followed by toothrow and then molar length. Higher *R*^2^ values indicate stronger correlations between the independent and dependent variables. Thus, skull length has the strongest predictive power, followed by toothrow length. Molar length has the weakest relationship.

| Measurement | Slope | Intercept | R<sup>2</sup> | p-value |
| --- | --- | --- | --- | --- |
| Skull Length (mm) | 3.759 | -3.793 | 0.97 | <0.001 |
| Lower Right Toothrow Length (mm) | 2.906 | -0.438 | 0.91 | <0.001 |
| Lower Right first molar length (mm) | 2.720 | 0.661 | 0.79 | <0.001 |

**Table 4.** Reduced major axis regressions for the relationship between skull and dental characters and body mass in Arvicolinae and Sigmotondinae.

| Measurement | Slope | Intercept | R<sup>2</sup> | p-value |
| --- | --- | --- | --- | --- |
| Skull Length (mm) - Arvicolinae | 3.81 | -3.85 | 0.98 | <0.001 |
| Lower Right Toothrow Length (mm) - Arvicolinae | 3.78 | -1.44 | 0.97 | <0.001 |
| Lower Right first molar length (mm) - Arvicolinae | 3.71 | -0.32 | 0.97 | <0.0001 |
| Skull Length (mm) - Sigmodontinae | 3.69 | -3.70 | 0.96 | <0.001 |
| Lower Right Toothrow Length (mm) - Sigmodontinae | 3.02 | -0.44 | 0.95 | <0.001 |
| Lower Right first molar length (mm) - Sigmodontinae | 3.40 | 0.59 | 0.92 | <0.001 |

**Fig. 1.**
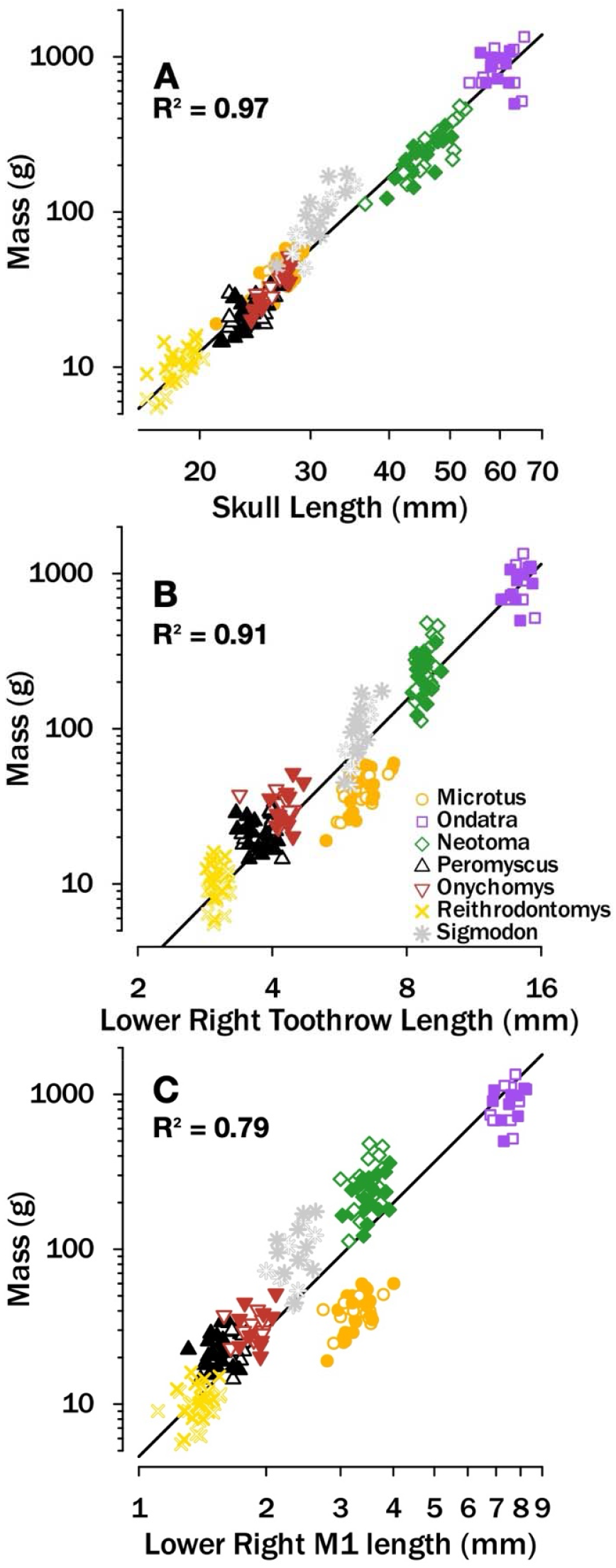
Body mass is significantly positively correlated with skull length (A), lower right toothrow length (B), and lower right first molar (M1) length (C) for species in family Cricetidae. Open symbols represent males, closed symbols represent females. Body mass and skull measurements were reported in log base 10.

**Fig. 2.**
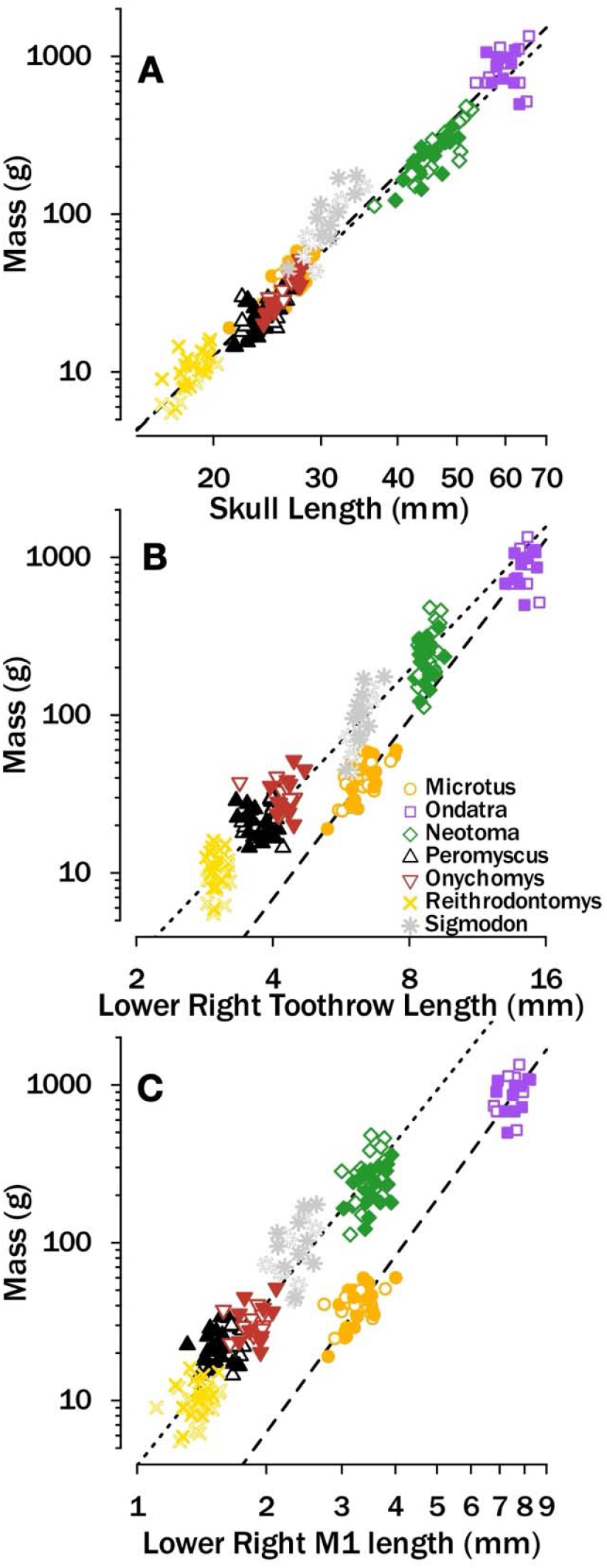
Body mass is significantly positively correlated with skull length (A), lower right toothrow length (B), and lower right first molar (M1) length (C) for each of the subfamilies. Open symbols represent males, closed symbols represent females. Dotted lines are for the Sigmodontinae. Dashed lines are for the Avicolinae. Body mass and skull measurements were reported in log base 10.

## Discussion

The aim of this study was to determine how skull length, toothrow length, and first molar length are related to body size in cricetid rodents from Nebraska. I found a significant positive relationship between all skull and dental characters and body size for rodents of the family Cricetidae from Nebraska (Fig. 1, Table 3). Individuals of larger body size have longer skulls, toothrows, and molars. Other studies have shown that skull and dental characters like molar length, skull length, and toothrow length are significantly related to body size in other groups of mammals (Bertrand et al. 2015, Freudenthal and Martín-Suárez 2013, Damuth and McFaddent 1990). Therefore, it is not surprising that larger cricetid rodents have larger skull and dental characters. Because the reduced major axis regressions for each skull and dental character are significant, any of the measured skull and dental characters can be used to predict body size in cricetid rodents.

An additional aim of this study was to determine which of the measured skull and dental characters were the best at predicting body size in cricetid rodents. Skull length is the best character to use in determining body size as it has the strongest correlation with the highest *R*^2^ value. Lower first molar length is the worst character to use when attempting to predict body size of the three measured, as it has the weakest correlation with the lowest *R*^2^ value. Toothrow length also makes for a good predictor for body size as it has an *R*^2^ value of above 0.9 (Table 3). Lower first molar length is not as good of a predictor due to the fact that one genus (*Microtus*) of Cricetidae has an unusually large lower first molar length for its body size (Fig. 1C). The genus *Microtus* is in the subfamily Arvicolinae, whereas all other genera but one (*Ondatra*) are in the subfamily Sigmodontinae. It appears that arvicolines have larger lower first molar lengths than other Cricetidae at a given body size. Other studies that have focused solely on the arvicolines found a much steeper relationship between lower first molar length and body size (Martin 1993, Martin, et al. 2009).

A final aim of this study was to determine if the relationship between skull and dental characters and body size was the same for the subfamilies Arvicolinae and Sigmodontinae. The relationship for skull length was similar for arvicolines and sigmodontids (Fig. 2, Table 4). However, the relationships for lower first molar length and toothrow length with body size were different for the two subfamilies (Fig 2B, C, Table 4). The arvicolines have a much steeper slope for the relationship between toothrow length and body size than the sigmodontids. In addition, the intercepts of the relationships are offset (Fig. 2B, Table 4). The slope of the relationship between lower M1 length and body size is similar arvicolines and sigmodontids, but the intercepts are offset. The regression equation between M1 length and body size found here is similar to the one found by Martin (1993) despite a larger sample size in this study and the geographic restriction of specimens to Nebraska. Importantly, deriving regression equations for the arvicolines and sigmodontids separately provides much stronger predictive power than when analysed together. When possible, regression equations for the individual subfamilies should be preferentially used.

The overall sample size was large for this kind of study (Betrand et al. 2015, Martin 1993, Freudenthal and Martín-Suárez 2013, Hopkins 2008, 2018), and included 10 males and 10 females from all but one species, which is enough to characterize morphological variation in a population (see Smith, et al. 1998). However, the species measured are only ones that occur in Nebraska and all specimens were from Nebraska. There are over 800 species of Cricetidae (Wilson and Reeder 2005), of which this study measured 11. Although it is encouraging that the relationship between M1 length and body size for arvicolines found in this study is similar to that of Martin (1993), it is possible that the relationships between skull and dental characters and body size will be different if more species are added. There are a number of ways to expand upon this study including, but not limited to, asking whether the relationships between skull and dental characters and body size in North American Cricetidae are similar to those on other continents. In particular, the species richness of cricetids in Asia is high (Wilson and Reeder 2005). Therefore, comparing relationships in North America to those of Asia would be insightful in how to best predict body size using the skull and dental characters of skull length, toothrow length, and lower first molar length. This study itself only contains one family of rodents, Cricetidae. This could be expanded to include other families within the order Rodentia to see if the results are similar in other families.

## Conclusion

This study was able to distinguish which measured skull and dental character was most accurate for predicting body size in cricetid rodents. For the family Cricetidae, skull length and toothrow length are the best predictors of body size (Fig. 1, Table 3). M1 length is a poorer predictor. However, when separated by subfamily, M1 length is as good a predictor of body size as skull length and toothrow length (Fig. 2, Table 4). Measuring three skull and dental characters also provides more options for palaeontologists to use when estimating body size in extinct rodents given the scrappy nature of the fossil record (Behrensmeyer 1984). Future work should expand the number of species included to get a more robust estimate for the relationship between body size and skull and dental characters for Cricetidae. Moreover, future work should include species from other continents to determine if the scaling relationships between skull and dental characters and body size in Cricetidae vary across the globe.

## Acknowledgements

I thank Bob Zink and Rob Wilson of the University of Nebraska State Museum for access to specimens. I thank Peter Wagner for help with data analyses and for comments on a previous version of the manuscript.

